## Supplemental Figure 1 for "Kynurenic acid, a key L-tryptophan-derived metabolite, protects the heart from an ischemic damage"

### Slide 1
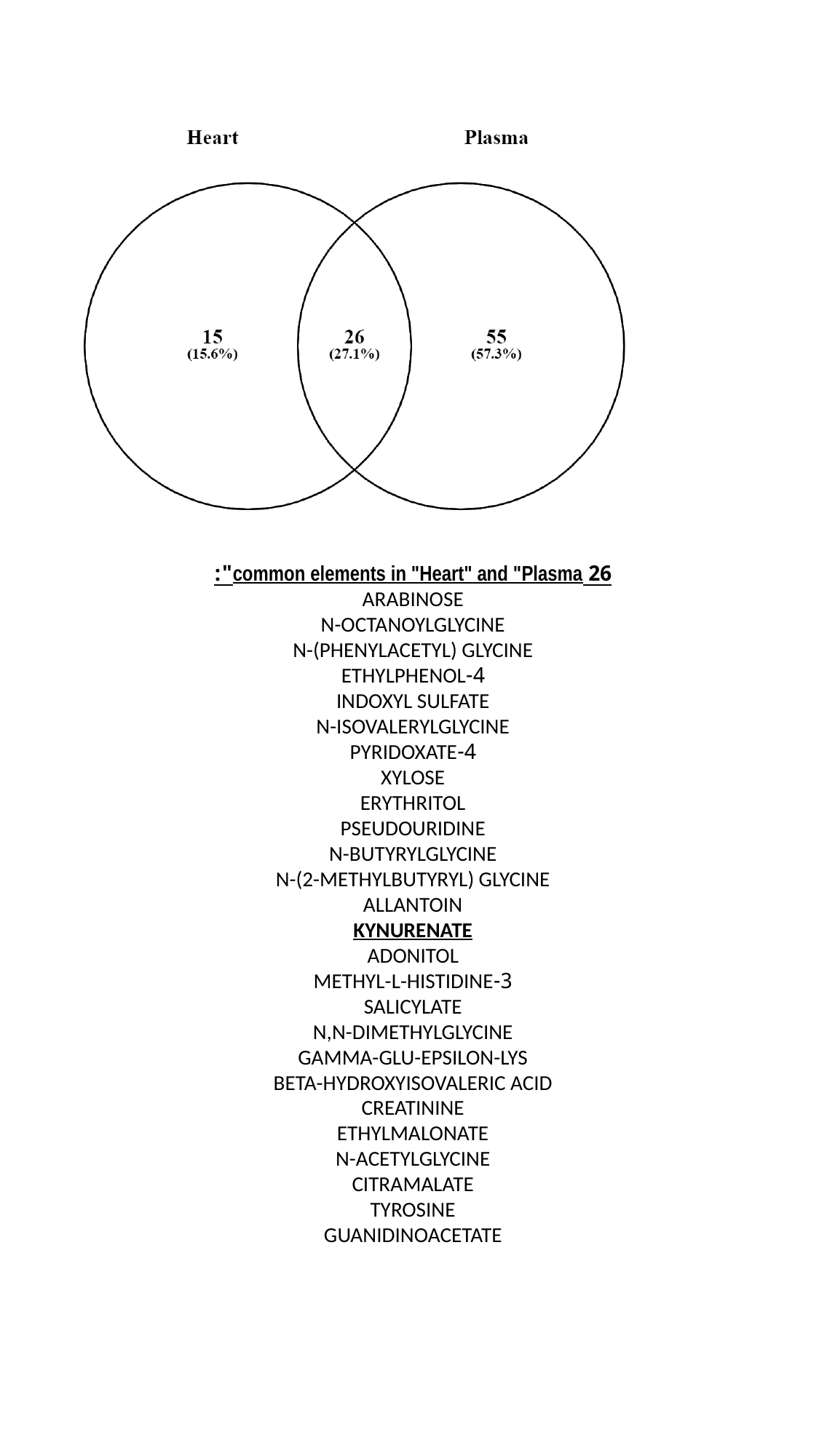

26 common elements in "Heart" and "Plasma":
ARABINOSE
N-OCTANOYLGLYCINE
N-(PHENYLACETYL) GLYCINE
4-ETHYLPHENOL
INDOXYL SULFATE
N-ISOVALERYLGLYCINE
4-PYRIDOXATE
XYLOSE
ERYTHRITOL
PSEUDOURIDINE
N-BUTYRYLGLYCINE
N-(2-METHYLBUTYRYL) GLYCINE
ALLANTOIN
KYNURENATE
ADONITOL
3-METHYL-L-HISTIDINE
SALICYLATE
N,N-DIMETHYLGLYCINE
GAMMA-GLU-EPSILON-LYS
BETA-HYDROXYISOVALERIC ACID
CREATININE
ETHYLMALONATE
N-ACETYLGLYCINE
CITRAMALATE
TYROSINE
GUANIDINOACETATE
