## Supplemental Figure 2 for "Kynurenic acid, a key L-tryptophan-derived metabolite, protects the heart from an ischemic damage"

### Slide 1
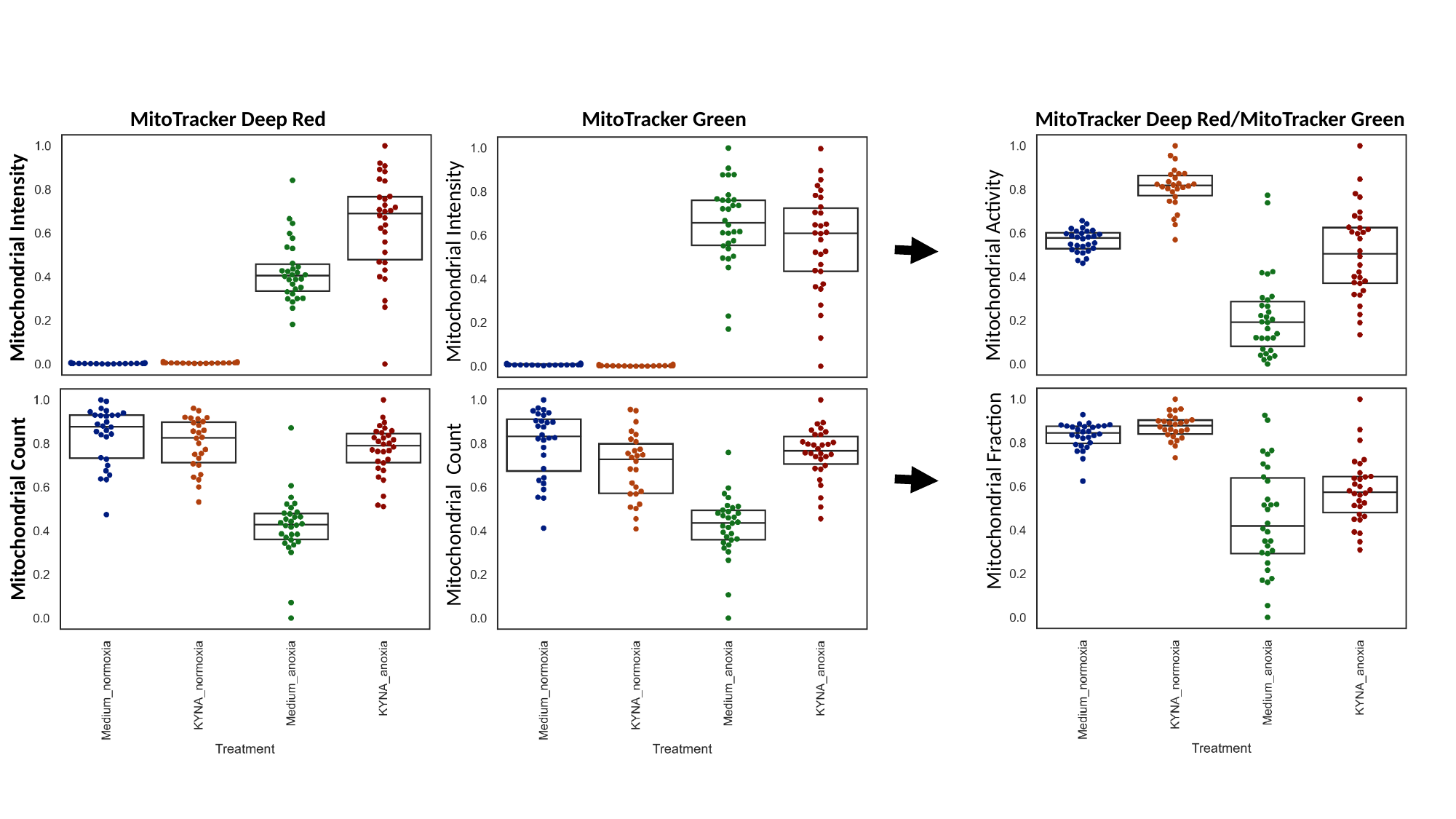

MitoTracker Deep Red
MitoTracker Green
MitoTracker Deep Red/MitoTracker Green
Mitochondrial Intensity
Mitochondrial Intensity
Mitochondrial Activity
Mitochondrial Fraction
Mitochondrial Count
Mitochondrial Count
